# Assessing the use of ellipsoidal microparticles for determining lipid membrane viscosity

**DOI:** 10.1101/2021.08.22.457294

**Authors:** P. E. Jahl, R. Parthasarathy

## Abstract

The viscosity of lipid membranes sets the timescales of membrane-associated flows and therefore influences the dynamics of a wide range of cellular processes. Techniques to measure membrane viscosity remain sparse, however, and reported measurements to date, even of similar systems, give viscosity values that span orders of magnitude. To address this, we improve a method based on measuring both the rotational and translational diffusion of membrane-anchored microparticles and apply this approach and one based on tracking the motion of phase-separated lipid domains to the same system of phase-separated giant vesicles. We find good agreement between the two methods, with inferred viscosities within a factor of two of each other. Our technique uses ellipsoidal microparticles, and we show that the extraction of physically meaningful viscosity values from their motion requires consideration of their anisotropic shape. The validation of our method on phase-separated membranes makes possible its application to other systems, which we demonstrate by measuring the viscosity of bilayers composed of lipids with different chain lengths ranging from 14 to 20 carbon atoms, revealing a very weak dependence of two-dimensional viscosity on lipid size. The experimental and analysis methods described here should be generally applicable to a variety of membrane systems, both reconstituted and cellular.

**Statement of Significance:** The lipid bilayers that underlie cellular membranes are two-dimensional fluids whose viscosity sets timescales of flow. Lipid membrane viscosity remains poorly quantified, with a paucity of methods and considerable disagreement between values reported using different techniques. We describe a method based on measuring the Brownian diffusion of ellipsoidal microparticles which we apply to phase-separated membranes alongside a previously established method for determining membrane viscosity, finding good agreement between the two techniques. We further examine homogenous membranes composed of lipids with different chain lengths, not amenable to phase-separation-based methods, revealing a very weak dependence of viscosity on lipid size. Our approach should be applicable to a wide range of membrane systems, both in vitro and in living cells.

## Introduction

Viscosity is a key determinant of lipid membrane behavior as it governs the force and time scales for the motion of membrane-embedded objects. Even for pure lipid bilayers, viscosity remains challenging to measure, as membranes are thin and fragile. In recent years several techniques have emerged, based in many cases on tracking the positions of membrane-associated objects including phase-separated lipid domains (1–6), microparticles or macromolecules anchored to membranes (7, 8), and microparticles near membranes (9). Nonetheless, considerable disagreement exists between reported membrane viscosity (η_m_) values. Even for bilayers composed almost completely of the simple phospholipid DOPC (1,2-dioleoyl-sn-glycero-3-phosphocholine), viscosity assessments at room temperature (21-25 °C) range from less than 0.6 × 10^−9^ Pa s m (10) to 64 ± 34 × 10^−9^ Pa s m (11). Different experiments on freestanding DOPC bilayers spanning apertures in solid supports give η_m_ < 0.6 × 10^−9^ Pa s m (10) and η_m_ = 16 ± 3 × 10^−9^ Pa s m (8), the former using optical traps to apply and assess forces on either side of the membrane, the latter analyzing the translational and rotational diffusion of membrane-anchored microsphere pairs. Measurements of domain diffusion in giant unilamellar vesicles (GUVs) with three major components including DOPC give less varied results, with η_m_ around 1-10 × 10^−9^ Pa s m (4, 6, 12).

It is unclear whether all these discrepancies result from flaws in methodologies, differences in membrane properties induced by different platforms and geometries, or other issues. Freestanding “black” lipid membranes, for example, are formed using solvents that initially separate two lipid monolayers and that must fully evaporate to give a bilayer, a condition that is often difficult to check. GUVs avoid this issue, but the use of phase separated domains to determine viscosity of course restricts their application to compositions that are capable of phase separation.

Applying two different viscosity characterization methods to the same type of lipid membrane would improve our understanding of the methods’ accuracy and applicability. We therefore examined the Brownian motion of phase separated domains in GUVs and examined the translational and rotational diffusion of microparticles anchored to similarly formed or identical GUVs. The phase separated vesicles make use of a well-known miscibility transition in ternary mixtures of cholesterol and lipids with saturated and unsaturated acyl chains; below a critical temperature, the bilayer separates into coexisting liquid ordered (“L_O_”) and liquid disordered (“L_D_”) phases (13, 14). The circular domains of the minority phase act as membrane inclusions of varied but well-defined and optically measurable radii. The domains’ Brownian trajectories can be analyzed by 1-point (1, 12) or 2-point (4) microrheological methods to reveal the underlying viscosity.

Analyzing the diffusion of membrane-anchored particles is, in principle, applicable to all lipid compositions. Knowing the translational diffusion coefficient D_T_ and the radius of the attachment *a*, one can infer the two-dimensional viscosity of the membrane via hydrodynamic models such as that of Hughes, Pailthorpe, and White (HPW) (15), a general extension of the large viscosity / low radius analysis of Saffman and Delbrück (16). However, *a* is not well known in practice; it may differ from the particle radius due to the smaller extent of attachment sites, giving a lower effective *a*, or membrane deformation, giving a larger effective *a* (8). For this reason, Hormel et al. developed the technique of linking two spherical particles together, one of which was coated with streptavidin proteins that bind to a small fraction of biotinylated lipid, the other of which, coated with biotin that binds to streptavidin on the first bead, serves solely to make the pair anisotropic (Figure 1a). Measuring the rotational and diffusion coefficient (D_R_) as well as D_T_ enables determination of η_m_ and *a*, as demonstrated for freestanding bilayers previously (8).

**Figure 1.**
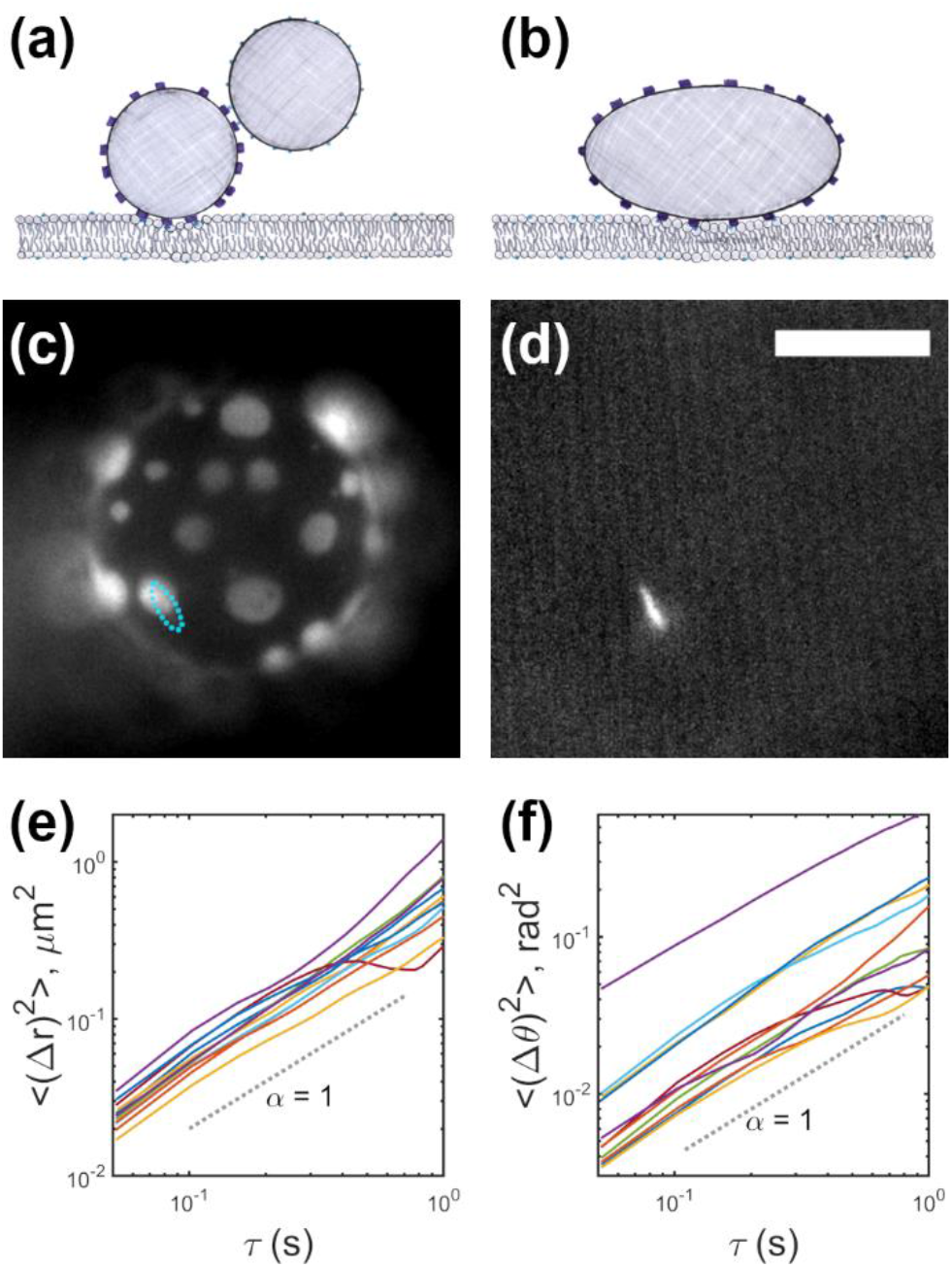
(a, b) Tracking the position and orientation of non-isotropic membrane-anchored particles allows determination of membrane viscosity and the effective size of the linkage. (a) Schematic of an earlier method, using pairs of spherical tracers. (b) Schematic of the method presented here, using single ellipsoidal tracers. (c, d) Simultaneously acquired fluorescence images of (c) a phase-separated giant unilamellar vesicle and (d) a membrane-anchored ellipsoidal particle. The particle position and orientation are indicated in (c) by the cyan oval. Scale bar: 10 μm. (e, f) Mean-squared displacements versus lag time for (e) translation and (f) rotation for vesicle-anchored particles. The dashed line, with slope 1, indicates Brownian diffusion.

To make this method easier to implement, we modify it here to use ellipsoidal particles, formed by stretching microspheres (17) and coating them with streptavidin (see Materials and Methods). This gives a non-circular attachment footprint (Figure 1b), assessed below, but simplifies sample preparation by avoiding delicate adjustment of concentrations to minimize formation of particle multimers or monomers.

We report here the comparison of membrane viscosity values derived from tracking ellipsoidal particles and phase separated domains in giant unilamellar vesicles, showing that if the ellipsoidal contact geometry is accounted for, the two methods are in good agreement. To illustrate the generality of the particle attachment method, we report the viscosity of a series of single component, non-phase-separating bilayers made of lipids with monounsaturated acyl chains of lengths ranging from 14 to 18 carbons, revealing a weak dependence of viscosity on chain length.

## Materials and Methods

### Giant Unilamellar Vesicles

GUVs were formed by electroformation (18) in 0.1 M sucrose. In brief, lipids of the desired composition dissolved in chloroform are deposited on glass coated with indium tin oxide (ITO) coated and allowed to dry under vacuum. The space between two such ITO-glass pieces, set by a silicone spacer, is filled with a sucrose solution. An alternating voltage across the ITO-glass leads to membrane swelling and the formation of GUVs.

### Lipid compositions

For experiments involving phase-separated GUVs, membranes were composed of 35.5% DPPC (1,2-dipalmitoyl-sn-glycero-3-phospocholine), 15.5% DOPC (1,2-dioleoyl-sn-glycero-3-phosphocholine), 40% cholesterol, 8% DOTAP (1,2-dioleoyl-3-trimethylammonium-propane (chloride salt)), 0.5% 16:0 biotinyl PE (1,2-dipalmitoyl-sn-glycero-3-phosphoethanolamine-N-(biotinyl) (sodium salt)), and 0.5% Texas Red DHPE (1,2-Dihexadecanoyl-sn-glycero-3-phosphoethanolamine, Triethylammonium Salt). Texas Red DHPE selectively partitions into the L_D_ phase of phase separated vesicles. DOTAP is an unsaturated cationic lipid whose presence, we find, increases the likelihood of particle binding, presumably due to the negative surface charge of polystyrene. The ratio of unsaturated : saturated (DPPC) : cholesterol lipids is therefore 23.5 : 35.5 : 40, which gives a majority L_O_ phase with circular L_D_ domains (Figure 1c).

For experiments on homogeneous vesicles, membranes were composed of 0.5% 16:0 biotinyl PE, 0.5% Texas Red DHPE, 8% DOTAP, and 91% of either 1,2-dimyristoleoyl-sn-glycero-3-phosphocholine (14:1 (Δ9-Cis) PC), 1,2-dipalmitoleoyl-sn-glycero-3-phosphocholine (16:1 (Δ9-Cis) PC), 1,2-dioleoyl-sn-glycero-3-phosphocholine (18:1 (Δ9-Cis) PC (DOPC)), and 1,2-dieicosenoyl-sn-glycero-3-phosphocholine (20:1 (Δ11-Cis) PC). The double bonds of the lipids with 14, 16, and 18 carbon chains all occur at the ninth carbon atom; the double bond of 20 carbon chain occurs at carbon 11.

### Ellipsoidal particles

Fluorescent, ellipsoidal polystyrene microparticles were formed based on the method described in Ref. (17). In brief, 1 micron diameter fluorophore-containing polystyrene microspheres (ThermoFisher Scientific “FluoSpheres,” catalog number F8776; excitation and emission peak wavelengths 505/515 nm) are placed in a solution of 6.2% polyvinyl alcohol and 2.5% glycerol by mass in deionized water. Evaporation leaves the particles embedded in a thin, flexible sheet. After liquification of the embedded particles by continuous immersion in toluene, the sheet is placed in a home-made mechanical stretching device, stretched, and removed from the toluene allowing the particles to solidify in the shape of the now deformed cavities. The film is then immersed in water, dissolved, and the microparticles are isolated by centrifugation. The resulting particles are prolate spheroids with major axis of 3.3 ± 0.6 μm (mean ± standard deviation) and minor axis 1.3 ± 0.3 μm, assessed by optical microscopy. Particles are coated non-specifically by streptavidin (Sigma-Aldrich) by incubation overnight in a 0.01 g/ml streptavidin / phosphate buffered saline solution, then centrifuged and sonicated in a bath sonicator to break up aggregates. Streptavidin-coated particles were incubated with GUVs containing biotinylated lipids for one hour at a sufficiently low concentration that the majority of vesicles had zero or one particle bound. Higher incubation concentrations, at which vesicles had more than one bound particle, led to particle aggregation.

### Fluorescence microscopy

Vesicles were placed in 0.1 M sucrose and imaged using a Nikon TE2000 inverted fluorescence microscope with a 60X oil immersion objective lens at room temperature (296 ± 1 K). Images were recorded using a Hamamatsu ORCA-Flash 4.0 V2 sCMOS camera at 20-33 frames per second. To simultaneously record two color channels, one of the lipid domains labeled with Texas Red and one for the fluorescent ellipsoidal beads, a Cairn OptoSplit emission image splitter was used.

### Image analysis

The analysis of phase separated domain positions was as is described in (4). Domains were identified using intensity thresholding and their centers were found by fitting Gaussian profiles using maximum likelihood estimation. The boundaries of domains were found with a bilateral filter and the total pixel area of each domain was used to determine its radius. Domain positions across frames linked into tracks with a nearest neighbor linking, and tracks of less than 100 consecutive frames were rejected. Ellipsoidal microspheres were detected by identifying the brightest single object in each frame in the appropriate fluorescence emission channel. Particle centers were found using the symmetry-based algorithm described in Ref. (19). The orientation of the particle is determined by calculating the covariance matrix of intensity in the neighborhood around the center. The accuracy of the positional and orientational localization was assessed using simulated images of ellipses that mimic the size and signal-to-noise ratio of the data. To create the simulated images, a high-resolution image of an ellipse was convolved with the detection pointspread function based on the emission wavelength and numerical aperture, pixelated, and subjected to Poisson-distributed noise. The localization accuracies were 35 nm in position and 0.0044 radians in angle, leading to overall uncertainties in diffusion coefficients and viscosities that are small compared to the variability across samples.

### Diffusion coefficients

From positions and angles, translational and rotational diffusion coefficients were calculated using the covariance-based method of Vestergaard et al. (20), which provides greater accuracy than linear fits of mean-square displacements and also provides estimates of localization accuracy and goodness of fit to a pure random walk.

### Hydrodynamic models

As noted in the main text, Hughes, Pailthorpe, and White (HPW) developed a model for the hydrodynamic drag of circular inclusions in membranes that is valid for arbitrary values of ε = 2 η_ext_ *a* / η_m_, where η_ext_ is the viscosity of the three-dimensional fluid in which the membrane is embedded, η_m_ is the membrane viscosity, and *a* is the inclusion radius (15). In our experiments, the external fluid is 0.1 M sucrose, for which η_ext_ =1.01 mPa s. The relationship between drag and other parameters is complex, involving several infinite series, and cannot be expressed in simple closed-form equations. We use the full HPW model, truncating series at 36 terms. Over a range of membrane viscosity, η_m_, and inclusion radius, *a*, we calculate D_T_ and D_R_ and determine the η_m_ and *a* that minimize their squared deviation from the measured D_T_ and D_R_.

To analyze the Brownian motion of rod-like particles, we use the hydrodynamic model of Levine, Liverpool, and MacKintosh (LLM) discussed in the main text (21). LLM evaluate the translational drag coefficients of displacements parallel (D_∥_) and perpendicular (D_⊥_) to the rod axis, and the rotational drag coefficient D_R_, all of which involve functions c(λ) of the dimensionless variable λ = 2 η_ext_ L / η_m_ (21). As with the HPW model, simple equations for c(λ) are unavailable. We interpolated c(λ) (see Equations 1–3) for rods with an aspect ratio of 3 based on published graphs spanning a large parameter range (21) (A. J. Levine and F. C. MacKintosh, personal communications). To determine membrane viscosity, we calculate D_∥_ and D_R_ over a range of membrane viscosity, η_m_, and ellipse major axis length, *L*, and find the η_m_ and *L* that minimize the squared deviation from the measured D_∥_ and D_R_. We sample over uncertainties in D_∥_ and D_R_ to determine the uncertainty in η_m_ and *L*, repeating the above process with 500 iterations over Gaussian distributions of D_∥_ and D_R_ with standard deviations equal to the D_∥_ and D_R_ uncertainties. The resulting η_m_ distribution is skewed; we report the half-width of its 68% confidence interval (equivalent to 1-σ for a Gaussian distribution) as the uncertainty in membrane viscosity. Final viscosity values for the full set of data from beads on phase-separated vesicles, or from each chainlength of homogenous vesicles, are reported as the weighted average of individual η_m_ values with uncertainty being the weighted standard deviation or weighted standard error of the mean (estimated as the weighted standard deviation divided by the square root of the number of points), as indicated in the main text.

### Software

Our MATLAB code to analyze diffusion coefficients and infer membrane viscosity using both the HPW and LLM models is publicly available on GitHub: https://github.com/rplab/HPW_MembraneDiffusion.

### Data availability

All trajectories of all membrane-anchored microparticles (position and angle), as well as inferred diffusion coefficients and viscosity values, are provided in CSV (comma-separated values) files included as Supplemental Data. Data from microparticles attached to phase-separated data are in “beadData_PhaseSepVesicles.csv,” and from microparticles attached to homogenous vesicles composed of lipids with different lengths are in “beadData_ChainLength_XX.csv,” where XX indicates the chain length.

## Results

We assessed the membrane viscosity of phase-separated GUVs (see Materials and Methods) by considering the Brownian motion of liquid-disordered domains and, separately, of membrane-attached fluorescent ellipsoidal microparticles. We first describe domain-derived viscosity values.

The minority phase of phase-separated lipid vesicles forms domains that behave as circular inclusions in a two-dimensional liquid (Figure 1B). A variety of experiments have verified that the hydrodynamic model of Hughes, Pailthorpe, and White (HPW) (15) links domain diffusion coefficients to domain radius and membrane viscosity (1, 3–5). Using the same methods as in prior work (Materials and Methods; (4, 5)), we imaged and tracked domain motion, determined diffusion coefficients, and related these to viscosity via the HPW model. From 13 GUVs composed of a ratio of 23.5 : 35.5 : 40 unsaturated-chain lipids : saturated-chain lipids : cholesterol (see Materials and Methods), each GUV having 2 to 18 L_D_ domains, we calculated an L_O_ phase viscosity of η_m_ = 1.8 ± 0.3 x 10^-9^ Pa s m (weighted mean ± weighted standard error of the mean, *N*=13). This is similar to the values reported in the literature for vesicles with a similar compositions, 20 : 40 : 40 saturated : unsaturated : cholesterol, namely η_m_ = 3.9 ± 0.4 x 10^-9^ Pa s m (4) and 1.9 ± 0.2 x 10^-9^ Pa s m (5).

We formed fluorescent ellipsoidal microparticles by stretching fluorescent polystyrene microspheres and anchored them to GUVs with a streptavidin / biotin linkage (see Materials and Methods). Imaging of domains and of microparticles were performed on the same batches of GUVs, and in some cases the same individual GUVs. Imaged GUVs had at most one bound microparticle.

Particle trajectories and diffusion coefficients were measured and calculated as described in Materials and Methods, using symmetry-based localization for particle position (19), moments of the intensity distribution for particle orientation, and a covariance-based estimator for diffusion coefficients that gives greater accuracy than fits to mean-squared-displacements (20). Nonetheless, it is informative to plot mean-squared-displacements, <Δr^2^(τ)> and <Δθ^2^(τ)> for position and angle, respectively, each which should scale with time lag τ as τ^α^ with α=1 for Brownian diffusion. We find α consistent with 1 as expected, with α = 1.1 ± 0.3 for translation and α = 0.9 ± 0.2 for rotation (mean ± std. deviation from *N* = 11 particles; Figure 1d, e).

We first assessed whether the Brownian dynamics of ellipsoidal microparticles is amenable to analysis with the HPW model, which considers circular membrane inclusions. This is not obvious a priori; the particles are clearly not circular, but the size and shape of the actual contact with the bilayer, or potential membrane deformations, is unknown. As in earlier work (8) we treated the membrane viscosity and an effective particle radius *a* as parameters of the HPW model. We show in Figure 2a contours in the D_T_-D_R_ plane for fixed η_m_; points along each contour correspond to different *a*. The upper left region corresponds to (D_T_, D_R_) values that are not physically realizable in the HPW model. We also plot in Figure 2a measured D_T_ and D_R_ values for GUV-anchored ellipsoidal particles, several of which lie in the un-physical regime. This suggests that the HPW model is inappropriate for this system, and the shape anisotropy of the particles is important.

**Figure 2.**
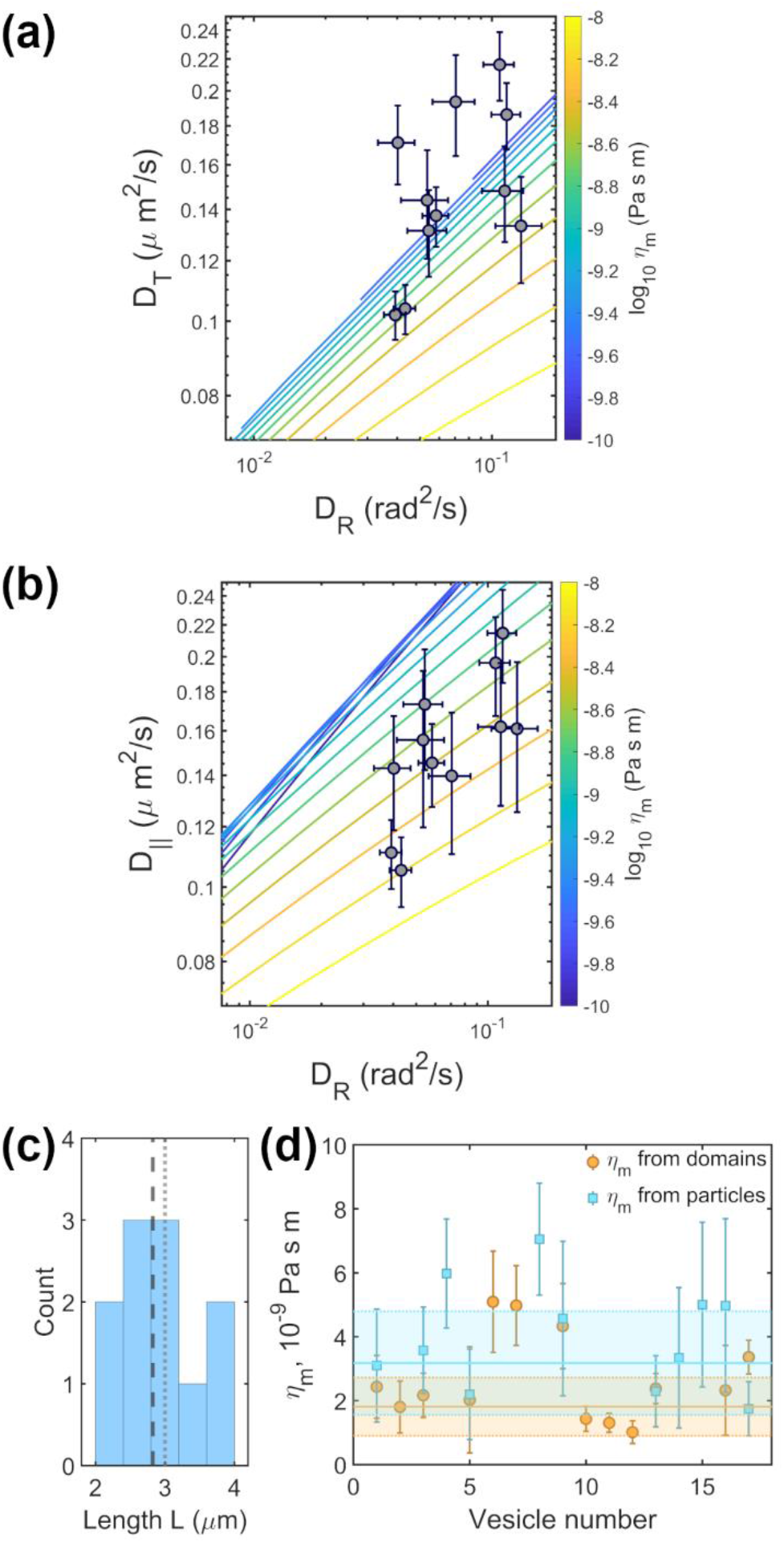
(a) Measured values of translational (D_T_) and rotational (D_R_) diffusion coefficients of membrane-anchored elliptical particles, together with equal-viscosity contours calculated for circular membrane inclusions. Several datapoints occupy the physically inaccessible region outside the range of the contours. (b) Measured values of D_R_ and diffusion coefficients parallel to the particle’s long axis (D_∥_), together with equal-viscosity contours calculated for rod-like membrane inclusions of aspect ratio 3. All datapoints occupy physically accessible regions of the parameter space. (c) Histogram of the effective rod lengths, *L*. The dashed line indicates the mean value, and the dotted line the mean rod length assessed from optical measurements. (d) All membrane viscosity values derived from 17 GUVs, either from domain motion or ellipsoidal particle motion. Mean values and standard deviations are indicated by solid lines and colored bands, respectively.

Levine, Liverpool, and MacKintosh (LLM) developed a hydrodynamic model for extended, rodlike inclusions in two-dimensional fluids (21), validated by experiments on liquid crystal films (22). LLM analyzed hydrodynamic drag for translation parallel and perpendicular to the rod axis, as well as the rotational drag, yielding diffusion coefficients that can be written as:

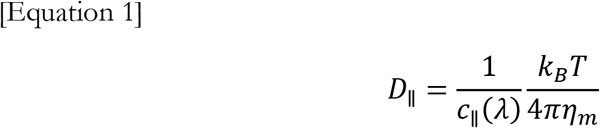

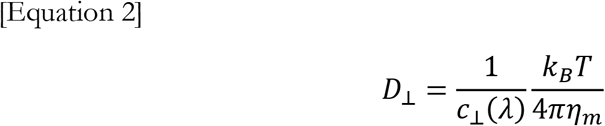

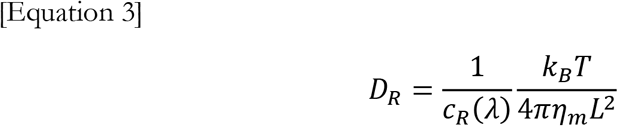

for translational motion parallel to the rod axis, translational motion perpendicular to the rod axis, and rotational motion, respectively. Here, *L* is the rod length, k_B_ is Boltzmann’s constant, *T* is the temperature, and the c(λ) are functions of the dimensionless parameter λ = 2 η_m_ *L* / η_m_. To apply the LLM model to membrane-anchored elliptical microparticles, we decompose frame-to-frame displacements into components parallel and perpendicular to the long axis of the ellipsoid and calculate the corresponding diffusion coefficients, D_∥_ and D_⊥_. The functions c_∥_(λ) and c_⊥_(λ) are very similar to each other for the range of λ probed in our experiments, and the determination of D_⊥_ is noisier than D_∥_ presumably due to worse localization along the thinner dimension and correlated uncertainty in rotational motion and motion perpendicular to the long axis. We therefore make use of D_∥_ and D_R_ only, determining the η_m_ and *L* for which diffusion coefficients calculated from the LLM model best-fit the observed values; see Materials and Methods. We note that *L* is treated as a fit parameter, since the effective length of the membrane contact may differ from the bead length.

Like the HPW-based analysis, we can plot for the LLM model equal-viscosity contours in the D_∥_-D_R_ plane, with points along each contour corresponding to different *L* (Figure 2b). Superimposing the measured D_∥_ and D_R_ values, we find that all datapoints are in the LLM model’s physically realizable regime (Figure 2b), implying that anisotropy is a significant factor in particle diffusion. The best-fit viscosity value is η_m_ = 3.2 ± 0.5 x 10^-9^ Pa s m (weighted mean ± weighted standard error of the mean, *N*=11).

The effective particle lengths *L* ranged from 2.5 to 3.4 μm (Figure 2c), with a mean ± standard error of 2.8 ± 0.1 μm, very similar to the physical length of 3.0 ± 0.3 μm.

We plot in Figure 2d all of the membrane viscosity values from 17 phase-separated GUVs, 7 of which featured both trackable domains and attached particles, along with the mean and standard deviation of each set of points as an indicator of the spread. For the 7 pairs of datapoints each from the same vesicle, η_m_ = 2.7 ± 0.2 x 10^-9^ Pa s m from domain data, and η_m_ = 2.5 ± 0.3 x 10^-9^ Pa s m from particle data (weighted ± mean standard error of the mean).

Overall, the close agreement of the η_m_ values derived from domain motion and ellipsoidal particle motion suggests that analysis of anisotropic particle diffusion provides an accurate measure of membrane viscosity, allowing it to be applied to other membrane systems. To illustrate, we next considered optically homogeneous membranes composed primarily (91%, see Materials and Methods) of one phosphatidylcholine lipid with one mono-unsaturated acyl chain. We varied the acyl chain length of this component, using four different lipids with 14, 16, 18, and 20 carbon chains; see Materials and Methods for chemical names and details of compositions. As above, fluorescent ellipsoidal microparticles were bound to GUVs, tracked, and analyzed using the LLM model.

The resulting membrane viscosities are plotted in Figure 3 for every examined GUV, along with averages for each composition. Viscosity values for the *C* = 14, 16, 18, and 20-carbon chain lipids were 1.6 ± 1.0, 2.0 ± 1.0, 1.9 ± 1.4, and 2.3 ± 1.1 x 10^-9^ Pa s m, respectively (weighted mean ± standard deviation). The values show a very weak increase of membrane viscosity with chain length, essentially indistinguishable from zero; a linear fit gives a slope dη_m_ /d*C* = 0.1 ± 0.2 x 10^-9^ Pa s m / atom (Figure 3).

**Figure 3.**
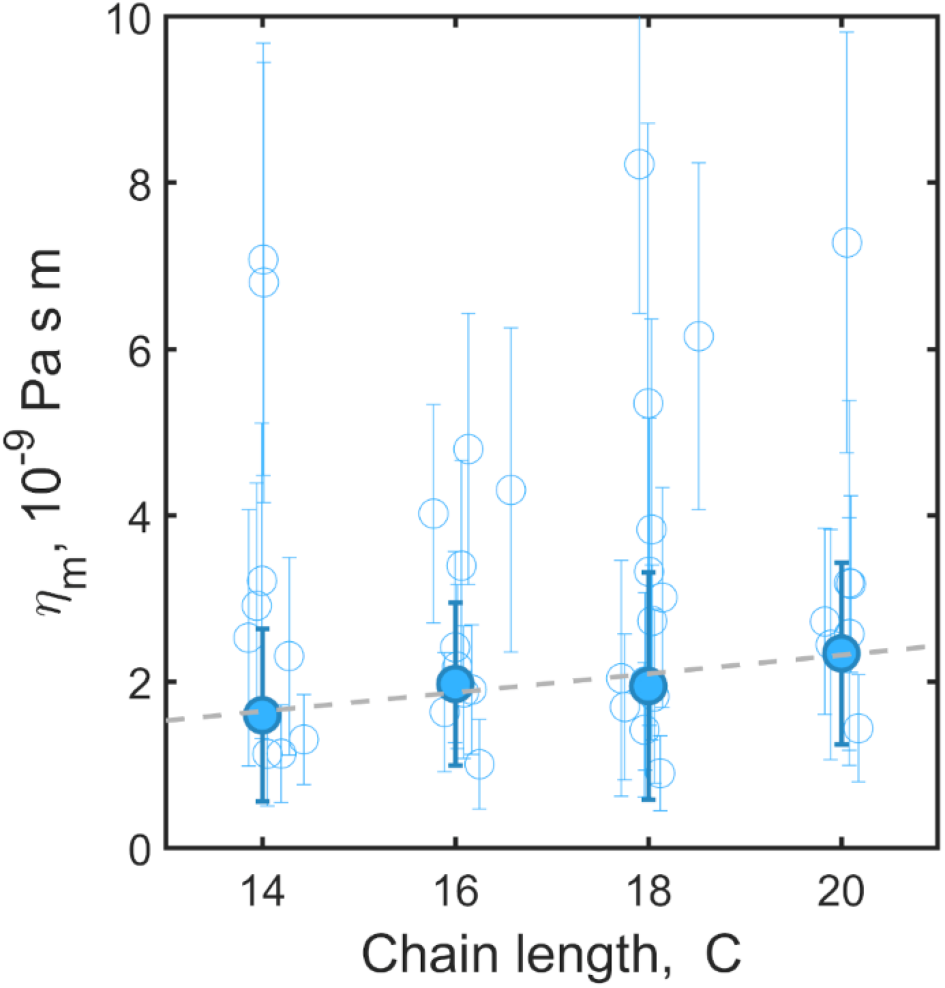
Viscosity of membranes as a function of lipid chain length. Plotted are values derived from the diffusion of individual membrane-anchored ellipsoidal particles (open circles) along with weighted mean and 68% confidence intervals (solid circles, error bars). The dashed line is a linear fit, with slope 0.1 ± 0.2 x 10^-9^ Pa s m / atom.

## Discussion

We have introduced a method for measuring the hydrodynamic viscosity of membranes using easily formed ellipsoidal microparticles. Techniques for quantifying viscosity are few in number, and measurements to date give orders-of-magnitude discrepancies for nominally similar lipid membranes. We validated our approach by investigating the same system, phase-separated three-component giant vesicles, with two different methods: assessment of the translational Brownian motion of domains and assessment of the translational and rotational Brownian motion of membrane-anchored ellipsoidal particles. Provided that the ellipsoidal geometry of the particles was accounted for, the two methods were in good agreement, within a factor of 2 of each other. We suggest, therefore, that ellipsoidal particles can be used quite generally as probes for membrane viscosity. Notably, the experimental uncertainties evident in this approach are large (Figure 2). However, we believe there is considerable room for improvement. For example, varying the particle size will change the magnitude of diffusion coefficients as well as the localization precision; there is likely some optimum, not explored here, that minimizes the uncertainty of the inferred viscosity.

We applied this ellipsoidal particle method to characterizing the viscosity of homogeneous vesicles composed of lipids with varying acyl chain lengths, which would be impossible to measure with techniques requiring phase separation. We found a very weak dependence of viscosity on chain length (Figure 3). It is interesting to compare the chain length dependence of η_m_ reported here with the chain length dependence of molecular diffusion. We caution that microparticle motion reflects a macroscopic hydrodynamic viscosity that may not apply at molecular scales, and it has long been noted that hydrodynamic models are of questionable utility for explaining molecular diffusion coefficients (23). Though systematic studies of lipid diffusion for a range of chain lengths are rare, Ramadurai et al. reported translational diffusion coefficient values measured using fluorescence correlation spectroscopy for the dye DiD in membranes composed of lipids with, as in this study, one double bond: 12.5, 9.6, 8.7, and 6.8 μm^2^/s for *C* = 14, 16, 18, and 20, respectively (24). Naively applying the continuum models of Hughes, Pailthorpe, and White or Saffman and Delbrück, using a molecular radius of 0.45 nm, gives viscosities of 0.14, 0.19, 0.21, and 0.29 x 10^-9^ Pa s m, respectively, about ten times lower than those inferred by studies of micron-scale domains or particles such as reported here (Figure 3). Intriguingly, the slope dη_m_/d*C* = 0.024 ± 0.003 x 10^-9^ Pa s m is also about one order of magnitude lower than what we derived from ellipsoidal particle data, suggesting a similar relative contribution of each additional carbon atom to both molecular-scale and macroscopic dissipation. We suggest further studies of membrane viscosity for membranes with different compositions, and at different temperatures, to better elaborate the fundamental connection between molecular structure and hydrodynamic response, which has remained opaque despite decades of interest.

The three-dimensional counterparts to the mono-unsaturated acyl chains of the lipids examined here (Figure 3) are alkenes, whose viscosities increase by about 200%, from η = 1.9 to 5.6 cP, over the range *C* = 14 (1-tetradecene) to *C* = 20 (1-eicosene) (25). Considering a thin film of alkenes equal in size to the thickness, *h*, of the lipid bilayers examined here (26) and naively assessing an effective two-dimensional viscosity as η_2D_ = η *h* gives an even stronger dependence of viscosity on chain length, increasing by about 300% from *C* = 14 to 20. The contrast with the very weak chain length dependence of lipid bilayer viscosity seen here highlights the fundamental importance of dimensionality in shaping hydrodynamic response; though not surprising, it is nonetheless remarkable.

Finally, we note that the ellipsoidal particle method we introduce here should be applicable to more complex, and more active, membranes, such as those of living cells. Though we made use of biotinylated lipids for particle attachments, one could just as well employ intrinsic cell-surface markers, and the extraction of the effective particle size, *L*, makes analysis independent of assumptions about the extent of linkage. We suspect that quantitative exploration of macroscopic membrane viscosity in a variety of systems will reveal unsuspected ways in which living systems modulate their hydrodynamic character.

## Supporting information

Supplemental Data (CSV files)

## Data and code availability

All trajectory data and inferred diffusion coefficients and viscosity values are provided as supplementary data, and MATLAB analysis code is publicly available, as described in Materials and Methods.

## Author Contributions

P.E.J. and R.P. designed the research. P.E.J. performed experiments. P.E.J. and R.P. analyzed data and wrote the manuscript.

## Acknowledgements

This material is based in part upon work supported by the National Science Foundation under Award Number 1507115. Any opinions, findings, and conclusions or recommendations expressed in this material are those of the author(s) and do not necessarily reflect the views of the National Science Foundation.

